# *Leishmania* allelic selection during experimental sand fly infection correlates with mutational signatures of oxidative DNA damage

**DOI:** 10.1101/2022.05.02.490304

**Authors:** Giovanni Bussotti, Blaise Li, Pascale Pescher, Barbora Vojtkova, Isabelle Louradour, Katerina Pruzinova, Jovana Sadlova, Petr Volf, Gerald F. Späth

## Abstract

Trypanosomatid pathogens are transmitted by blood-feeding insects, causing devastating human infections. Survival of these parasites in their vertebrate and invertebrate hosts relies on their capacity to differentiate into distinct stages that are the result of a co-evolutionary process. These stages show in addition important phenotypic shifts that often impacts infection, affecting for example parasite pathogenicity, tissue tropism, or drug susceptibility. Despite their clinical relevance, the evolutionary mechanisms that allow for the selection of such adaptive phenotypes remain only poorly investigated. Here we use *Leishmania donovani* as a trypanosomatid model pathogen to shed first light on parasite evolutionary adaptation during experimental sand fly infection. Applying a comparative genomics approach on hamster- isolated amastigotes and derived promastigotes before (input) and after (output) infection of *Phlebotomus orientalis* revealed a strong bottleneck effect on the parasite population as judged by principal component and phylogenetic analyses of input and output parasite DNA sequences. Despite random genetic drift caused by the bottleneck effect, our analyses revealed various genomic signals that seem under positive selection given their convergence between independent biological replicates. While no significant fluctuations in gene copy number were revealed between input and output parasites, convergent selection was observed for karyotype, haplotype and allelic changes during sand fly infection. Our analyses further uncovered signature mutations of oxidative DNA damage in the output parasite genomes, suggesting that *Leishmania* suffers from oxidative stress inside the insect digestive tract. Our results propose a new model of *Leishmania* genomic adaptation during sand fly infection, where oxidative DNA damage and DNA repair processes drive haplotype and allelic selection. The experimental and computational framework presented here provides a useful blueprint to assess evolutionary adaptation of other eukaryotic pathogens inside their insect vectors, such as *Plasmodium* spp, *Trypanosoma brucei* and *Trypanosoma cruzi*.

## INTRODUCTION

Leishmaniases are vector-borne diseases that generate important human morbidity and mortality worldwide [1]. These immuno-pathologies are characterized by a variety of clinical symptoms, including self-healing skin ulcers, disfiguring mucocutaneous lesions, and fatal hepato-splenomegaly. The etiological agents of leishmaniases are trypanosomatid pathogens of the genus *Leishmania* that exploit two major hosts, (i) phlebotomine sand flies, where they proliferate extracellularly as motile, insect-stage parasites termed promastigotes, and (ii) vertebrate (mostly mammalian) hosts that are infected during a sand fly blood meal, where promastigotes develop into immotile, proliferating amastigotes inside acidified macrophage phagolysosomes.

Aside stage differentiation, *Leishmania* uses an evolutionary strategy to adapt and increase fitness following the Darwinian paradigm of mutation and selection of the fittest [2]. As all microbial pathogens, *Leishmania* is constantly under selection by fluctuations in the various host environments, e.g. changes in the sand fly microbiome, the host immunogenetic makeup or the presence of anti-parasitic drugs. Unlike other pathogens that largely rely on stochastic changes in gene expression to generate selectable phenotypes [3, 4], *Leishmania* generates genetic and phenotypic variability through its intrinsic genome instability [5–8]. Frequent changes in chromosome and gene copy number cause gene dosage-dependent changes in transcript abundance, which have been linked to experimentally induced drug resistance, changes in tropism, or fitness gain during culture adaptation [9–15]. Aside this quantitative aspect on gene expression, we recently provided first evidence that gene dosage changes also allow for qualitative transcriptomic changes via haplotype selection, with chromosomal amplification being non-random as only observed for one of the two different haplotypes [14]. This was observed both *in vitro* during culture adaptation of the Sudanese isolate *L. donovani* Ld1S, but also during hamster infection *in vivo*, where transient trisomies seem to allow selection of tissue-specific allelic profiles in parasites recovered from spleen.

In contrast to mammalian infection, studies on *Leishmania* genomic adaptation inside its sand fly insect host are scarce and largely limited to karyotypic analyses and Single Nucleotide Variants (SNVs) profiling to monitor genetic hybridization events [11, 16–19]. To date, no information is available on other forms of genomic adaptation during vector infection. Here we address this important open question conducting experimental sand fly infection with *bona fide* amastigotes isolated from infected hamster spleen, and derived promastigotes with different karyotypic profiles. Applying our genome instability pipeline (GIP, [20]) on sand fly- recovered parasites revealed signature mutations of oxidative DNA damage that correlated with haplotype shuffling and allelic selection. These seem to drive parasite fitness gain during vector infection as judged by the convergence of allele frequency shifts across independent infection experiments.

## Material and Methods

### Animals and ethics statement

Two 7-8 weeks old female hamsters were used. Work on animals was performed in compliance with French and European regulations on care and protection of laboratory animals (EC Directive 2010/63, French Law 2013-118, February 6th, 2013). All animal experiments were approved by the Ethics Committee and the Animal welfare body of Institut Pasteur and by the Ministère de l’Enseignement Supérieur, de la Recherche et de l’Innovation (project n°#19683).

### Hamster infection and isolation of infectious amastigotes

Anesthetized hamsters were inoculated by intra-cardiac injection with 5 x 10^7^ infectious amastigotes obtained from infected hamster spleens (Figure 1). Hamster weight was monitored weekly and the animals were euthanized with CO_2_ before reaching the humane endpoint (20% weight loss). Spleens were collected and homogenized in PBS supplemented with 2.5 mg/ml saponine using the gentleMACS homogenizer with gentleMACS M tubes from Miltenyi Biotec. Amastigotes were purified as previously described [21] for DNA extraction, differentiation into promastigotes and further promastigote culture and sand fly feeding.

**Figure 1:**
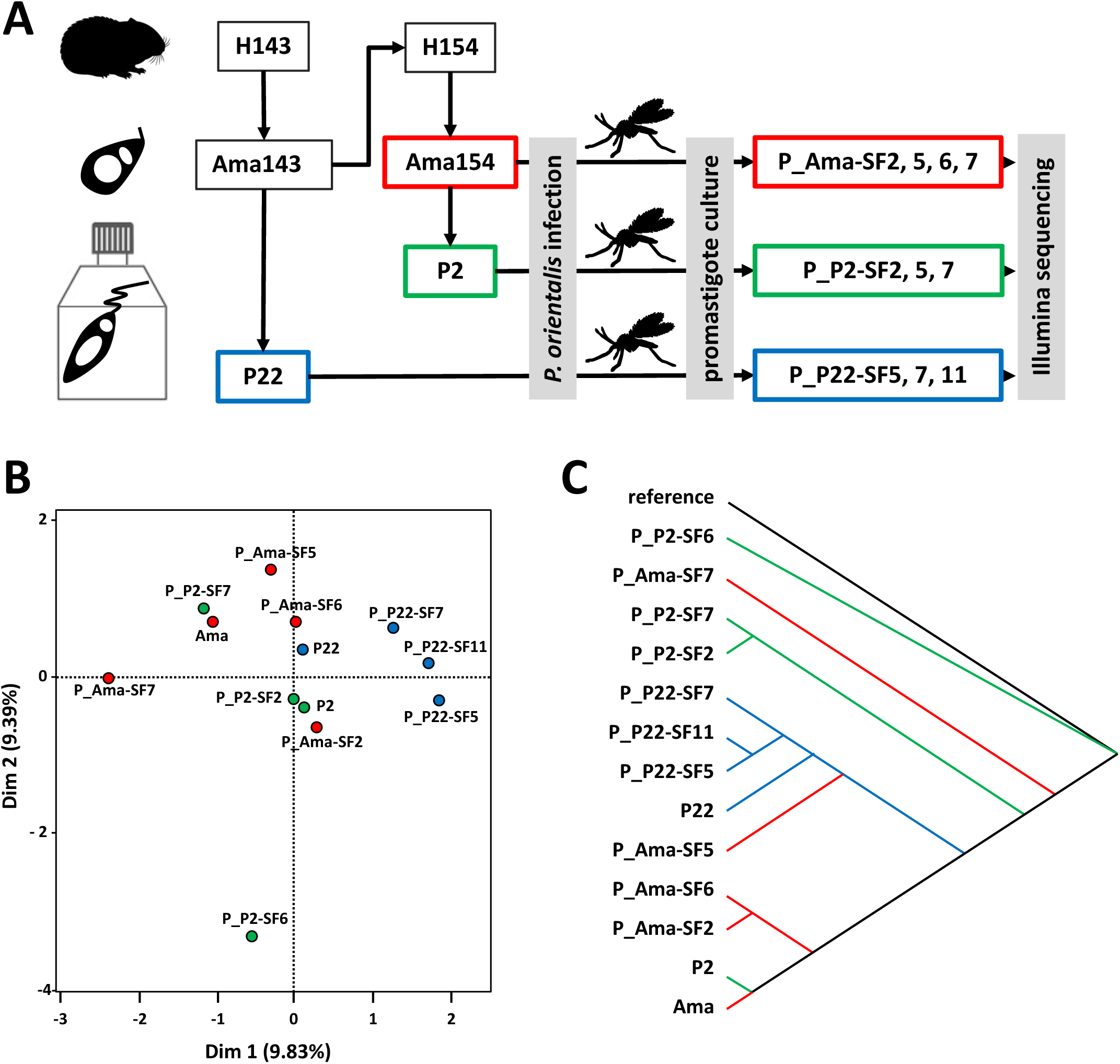
Experimental layout and phylogenetic analysis. (**A**) Overview of *L. donovani* Ld1S isolates origin before and after experimental sand fly (SF) infection. Splenic amastigotes (Ama) were obtained from infected hamsters (H) 143 and 154, adapted to promastigote culture for two (P2) and 22 (P22) passages, and the indicated parasites were fed to 100 *Phlebotomus orientalis* sand flies. Parasites were isolated 8 days post-infection and successful short-term cultures were established for 3 to 4 sand fly-derived promastigotes (SFn) (from a total of 10 dissected flies) per infection group. (B) Principal component analysis based on the average nucleotide identity scores between samples. The samples are represented by their projection along the two first component. (C) Phylogeny based on the SNVs of the samples. The tree is rooted on the Ld1S reference genome.

### Parasites and culture

*Leishmania donovani* strain 1S2D (MHOM/SD/62/1S-CL2D) was obtained from Henry Murray, Weill Cornell Medical College, New York, USA and maintained by serial passages in hamsters. Amastigotes were recovered from infected hamster spleen and differentiated into promastigotes in M199 complete medium (M199, 10% FBS, 25 mM HEPES; 100 µM adenine, 2 mM L-glutamine, 10 µg/ml folic acid, 13.7 µM hemin, 4.2 mM NaHCO_3_, 1xRPMI1640 vitamins, 8 µM 6-biopterin, 100 units penicillin and 100 µg/ml streptomycin, pH 7.4) at 26°C. Promastigotes were then maintained in culture by dilution in fresh medium once they are in stationary phase. At passage 2 (P2, Figure 1), parasites were either collected in exponential growth phase for sand fly feeding and DNA extraction or kept in culture until passage 22 (P22, Figure 1) and collected for DNA extraction and for sand fly feeding.

### Sand fly feeding and parasites recovery

Colonies of *P. orientalis* were maintained under standard conditions as previously described [22]. Three groups with 100 three to seven days old females were fed with 1 x 10^6^ parasites per ml suspensions of splenic amastigotes, promastigotes passage 2 or passage 22 respectively (Figure 1). After 8 days post-infection, 10 female sand flies were dissected, and parasites recovered from each sand fly. Promastigotes from individual sand fly were then expanded in culture (max. two passages) and collected at exponential growth phase for DNA extraction. Three to four promastigotes cultures of individually infected sand flies were derived from each of the three experimental groups (Ama, P2, P22).

### DNA extraction and deep sequencing

DNA extractions were performed using DNeasy blood and tissue kits from Qiagen according to manufacturer instructions. Nucleic acid concentrations were measured, and the DNA quality was evaluated on agarose gel. Sequencing was performed on DNA samples obtained before and after sand flies feeding. Short-insert paired-end libraries were prepared with the KAPA Hyper Prep Kit (Kapa Biosystems). The libraries were then sequenced using TruSeq SBS Kit v3-HS (Illumina Inc.), in paired end mode, 2 x 101bp, in a fraction of one sequencing lane of an HiSeq2000 flowcell v3 (Illumina Inc.) according to standard Illumina operation procedures at Centro Nacional de Análisis Genómico (CNAG).

### Genome sequencing and genome assembly data

The whole genome sequencing datasets described in this study were deposited in the Sequence Read Archive (SRA) [23] and available at the accessions reported in Table S1. All bioinformatics analyses were performed considering the *L. donovani* assembly GCA_002243465.1 of the 1S2D strain (https://www.ncbi.nlm.nih.gov/bioproject/PRJNA396645, BioSample SAMN07430226) and the annotations we previously published [10].

### Computational analyses

The read alignment, the quantification of chromosomes, genomic bins and genes, and the single nucleotide variant analyses were performed with the GIP pipeline and the giptools toolkit (version 1.0.9) [20]. The mapping statistics are provided in Table S2 and the GIP parameters are available in Dataset S1. Briefly, WGS reads were mapped by GIP using BWA-mem (version 0.7.17) [24]. The resulting alignment files were then sorted, indexed and reformatted by GIP using Samtools (version 1.8) [25]. Read duplicates were removed by GIP using Picard MarkDuplicates (http://broadinstitute.github.io/picard) (version 2.18.9). A minimum read alignment MAPQ score of 50 was enforced by GIP to select genes for cluster analysis and to call for SNVs. A full description of GIP pipeline steps and parameters are available from its online documentation (https://gip.readthedocs.io/en/latest/index.html).

GIP was used to compute the mean sequencing coverage of genomic bins (adjacent windows of 300bp) and genes. The somy scores shown in Figure 2A were computed based on the sequencing coverage of the genomic bins with mean read MAPQ greater than 50 (Tables S3-5). The bin coverage scores were normalized by the median genome coverage. The data was further scaled dividing by the median bin coverage of chromosome 25 — which is showing a stable disomic level across samples — and multiplied by two. In Figure 2B, gene coverage scores were normalized by median chromosome coverage to highlight amplifications or depletions with respect to the chromosome copy number (Table S6). In Figure S1 bin coverage scores were normalized by median genome coverage to account for differences in sequencing library size. GIP was used for the analysis of genes sharing high sequence identity, thus not directly quantifiable with the short-read technology adopted in this study. The pipeline grouped the nucleotide sequences of the genes with low mean MAPQ score into clusters with cd-hit-est (version 4.8.1) [26] with options ‘-s 0.9 -c 0.9 -r 0 -d 0 -g 1’. Next, for each cluster GIP computed the mean of the individual gene coverage scores normalized by median chromosome coverage.

**Figure 2:**
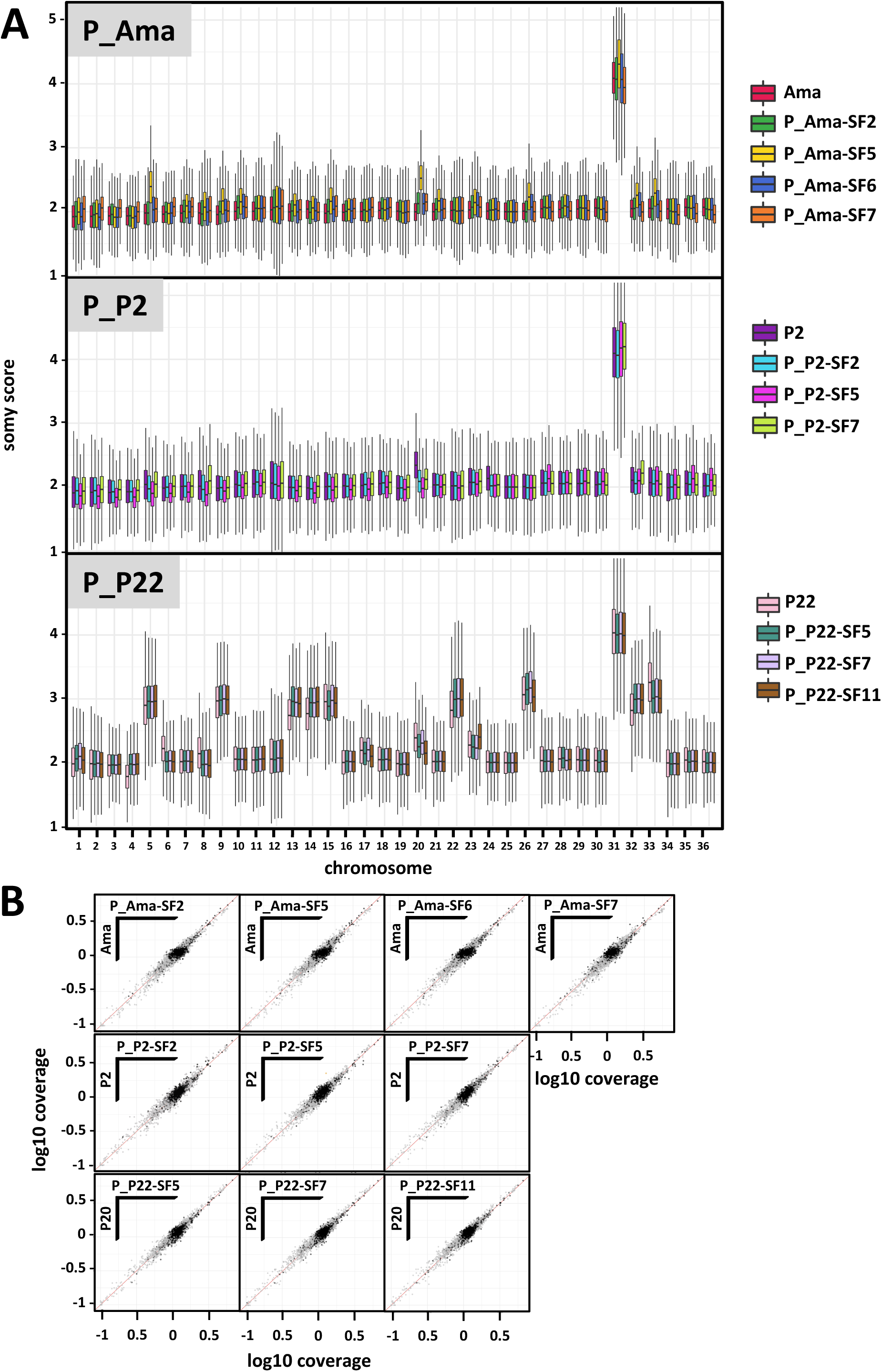
Copy number variation upon sand fly infection. (A) Chromosome ploidy analysis. The box plots represent the somy score distributions for each chromosome in the indicated samples. Outliers are not shown to ease plot readability. The boxes are colored according to the samples. (B) Gene copy number variation (CNV) analysis. In the scatter plots, individual genes are represented by dots. The plots show the log10 sequencing coverage of all genes as measured in the input (y-axes) and output strains (x-axes). The diagonal red lines represent the bisectors.

GIP was also used for the SNV analysis. The steps operated by GIP include SNV calling with Freebayes (version 1.3.2) followed by quality filtering accounting for both the frequency and the genomic context of the variant (see the pipeline documentation for more details). SnpEff (version 4.3t) [27] was used to predict the effect of the variant alleles on coding sequence. The predicted effects that were considered synonymous mutations are: ‘synonymous_variant’, ‘stop_retained_variant’ and ‘start_retained’. The predicted effects that were considered non synonymous mutations are: ‘missense_variant’, ‘start_lost’, ‘stop_gained’, ‘stop_lost’ and ‘coding_sequence_variant’. The phylogenetic tree was computed by the giptools using IQtree2 (version 2.1.2) with options ‘–seqtype DNA –alrt 1000 -B 1000’. To infer the tree GIP considered the set of filtered SNV excluding variants with frequency < 10% and adopting the IUPAC ambiguous notation for the positions with frequency ≤ 70%. Principal component analysis (PCA) was computed using giptools’ genomeDistance module, where pairwise distances between samples are based on their average nucleotide identity scores determined using the MUMmer package and PCA is preformed using the FactoMineR R package (version 2.3).

Further SNV analyses were performed based on the outputs of GIP, using custom Python 3.9 code relying on the following libraries: pysam [https://doi.org/10.1093/gigascience/giab007] (version 0.18), pandas [https://doi.org/10.25080/Majora-92bf1922-00a] (version 1.3.5), upsetplot [https://doi.org/10.1109/TVCG.2014.2346248] (version 0.6.0), matplotlib [https://doi.org/10.1109/MCSE.2007.55] (version 3.3.2), seaborn [https://doi.org/10.21105/joss.03021] (version 0.11.2) and biotite [https://doi.org/10.1186/s12859-018-2367-z] (version 0.31). Detection of loci with strong allele frequency changes between “input” and “output” samples relied on the following procedure. In the outputs of GIP, SNVs are characterized by a “ref” (reference) and an “alt” (alternative) allele, where the “ref” allele is the one present in the reference genome provided to GIP, and by the frequency of the “alt” allele in a sample. First, for a given “output” sample, a frequency “delta” was computed, as the difference between the alt allele frequencies in the output and in the input (a positive delta meaning an increase in the frequency of the “alt” allele in the output with respect to the input) (Tables S7-9). Strong allele frequency changes can therefore consist in a strongly positive delta, or a strongly negative delta. The absolute value of delta was computed (abs_delta), and its minimum across output samples of an experiment (min_abs_delta) was used to characterize a locus. The 95^th^ percentile of the distribution of min_abs_delta was used as a threshold above which a locus was considered as displaying strong allele frequency shifts (possibly convergent or divergent) (Figure S8). These loci were used to build “rainbow plots” such as the examples shown in Figure 5, and the same threshold was used to select the loci counted in the right heatmap of Figure 6D (for a given sample, a locus is counted if its abs_delta is above the 95^th^ percentile of the min_abs_delta distribution).

## RESULTS

### Experimental sand fly infection affects the genetic structure of the parasite population

*L. donovani* Ld1S splenic amastigotes (Ama) obtained from infected hamsters and derived promastigotes at early (passage (P) 2) and late (P22) stages of culture adaptation were fed to *Phlebotomus orientalis* sand flies (Figure 1A). Eight days after infection, 3-4 promastigote populations were isolated from individual flies for each experimental group of sand flies and expanded in culture for a maximum of two passages, which we previously showed is too short to have a significant impact on genomic adaptation [14, 28]. Following whole genome sequencing analysis, we first assessed the genomic distance between the various parental input parasites (Ama, P2, P22) and corresponding promastigote (P) sand fly (SF) output isolates (P_Ama-SF2, 5, 6, 7; P_P2-SF2, 5, 7; P_P22-SF5, 7, 11). Surprisingly, principal component analysis (PCA) based on average nucleotide identity (ANI) did not reveal defined clustering of corresponding input and output parasites, suggesting that sand fly infection significantly changes the genetic structure of the parasite population (Figure 1B). This result was confirmed by phylogenetic analysis based on SNVs, which revealed a polyphyletic organization for the Ama and P2 input and P_Ama-SF and P_P2-SF output parasites (Figure 1C). In contrast, P22 input and P_P22-SF output parasites clustered together, establishing their clear phylogenetic relationship. Our results are best explained by a strong bottleneck effect caused by sand fly infection [29–31], which reduces the genetic complexity of the P_Ama-SF and P_P2-SF output parasites compared to the genetically more heterogenous input strains, thereby affecting population structure and allele frequencies. Such bottleneck effects may be less pronounced in the P_P22-SF isolates, given that long-term culture of the P22 input parasites already reduced genetic heterogeneity by selecting for common aneuploidies and haplotypes that are beneficial for in vitro growth [14]. In the following, we investigated parasite genomic signals that arise during sand fly infection and tried to distinguish between their random nature due to the bottleneck effect or their non-random nature due to natural selection by assessing their reproducibility in independent biological replicates.

### Karyotypic changes during sand fly infection

We previously showed that copy number variations of both chromosomes and genes play an important role during promastigote fitness gain in culture [10, 14, 28]. Here we assess the link of these forms of genome instability to promastigote differentiation, proliferation and genetic adaptation during sand fly infection. We applied read depth analysis on the genome sequences of the various input and corresponding output parasites (see Figure 1) using our Genome Instability Pipeline (GIP) and the giptools analysis package [20]. The chromosomes of the Ama input parasites were largely disomic as judged by their somy score of two, with the exception of the stably tetrasomic chromosome (chr) 31 [6] that showed the expected somy score of four (Figure 2A, upper panel, Table S3). The sand fly-derived promastigote isolates P_Ama-SF2, 5, 6, 7 largely reproduced the karyotypic profile of their input Ama parasites, with the exception of chr 20 that is significantly amplified in P_Ama-SF5 showing a somy score of 2.5. This isolate further shows partial aneuploidy for chr 5, 9, 26, 32 and 33 as indicated by the increased median somy score. The absence of a similar amplification pattern in the other P_Ama-SF isolates suggests that this unique karyotypic profile is not under selection but rather result of the bottleneck event described in Figure 1. The karyotype of the P2 input parasites is largely disomic and thus resembles that of the Ama parasites, confirming our previous observations that only little karyotypic change is detected early during culture adaptation [14] (Figure 2A, middle panel, Table S4). Again, the derived SF isolates (P_P2-SF) largely reproduced the input karyotype as has been observed previously [11].

As expected from our previous experimental evolution studies, long-term culture adaptation of the P22 input parasites resulted in highly reproducible amplifications of chr 5, 9, and 26 [10, 14, 32] (Figure 2A, lower panel, Table S5). In addition to these driver aneuploidies, the P22 input parasites show an amplification of chr 6, 8, 13, 14, 20, 22, 32 and 33. Some of these chromosomes show significant somy score variations that converge in all three P_P22-SF output isolates, including increased copy number for chr 13, 14, 15, 22 and 32, or reduced copy number for chr 6, 8, 20, and 33. This convergence provides evidence for positive selection of beneficial and purifying selection against detrimental aneuploidies during P22 sand fly infection.

Similar to our karyotypic analysis, only minor changes were observed for gene Copy Number Variations (CNVs) between the Ama, P2 and P22 input parasites and their respective sand fly isolates (Figure 2B and S1, Table S6). For example, we observed gene amplification in individual SF isolates for the HSP83 gene cluster in P_AmaSF-5 (1,5-fold), for the snoRNA LD36Cs-1C2_c1084 in P_P2-SF5 (1,7-fold), and for the cluster of the hypothetical gene LdBPK_020011400 in P_P22-SF11 (1,5-fold). As they were only observed in individual samples, these CNVs don’t seem to be the result of selection but are likely to result from the bottleneck effect occurring during the sand fly infection.

Taken together, our analyses indicate that karyotypic selection can occur in the sand fly and may provide fitness advantage, at least in the P22 experimental group. Even though no karyotypic selection was observed for the Ama and P2 experimental groups under our experimental conditions, karyotypic adaptation may occur during prolonged promastigote proliferation following repeated blood feeding [33–35].

### Sand fly infection causes important changes in the SNVs profile

The read depth analyses above neither capture possible transient aneuploidies that may allow for haplotype selection, nor other forms of genome instability that may allow for haplotype or allelic selection. We next investigated these possibilities assessing the nature and frequency of single-nucleotide variants (SNVs). SNVs are substitutions of individual nucleotides at specific positions in the genome, which can give rise to gene variants (alleles) affecting gene function. SNVs are key drivers of evolutionary adaptation that can shape the phenotypic landscape through purifying and positive selection of respectively detrimental or beneficial alleles. Changes in SNV frequency in a population can be caused by bottleneck events, allelic selection, aneuploidy or recombination events. To distinguish between these possibilities, we investigated the SNVs distribution before and after sand fly infection and assessed the reproducibility of frequency changes across the independent SF isolates for a given experimental group.

We detected over 12,000 heterozygous SNVs across all input parasites (Ama, P2, P22) and their corresponding SF isolates compared to our recently generated reference genome (see Methods) (Figure 3A and S2, Tables S7-S9). While the majority of SNVs were shared between the various parental input and SF output parasites (>85%), a small number of SNVs were detected only in the three input parasites, which may have been lost in the output strains due to the bottleneck event described above (see Figure 1) that may cause loss of parasite sub- populations during sand fly infection. We also observed a number of new SNVs in the output strains, a significant number of which was shared between the biological replicates of a given experimental group (Figure 3A, red dots). The emergence of these new variants and their convergence in the output strains suggest that these SNVs may represent new mutations that are positively selected during sand fly infection.

**Figure 3:**
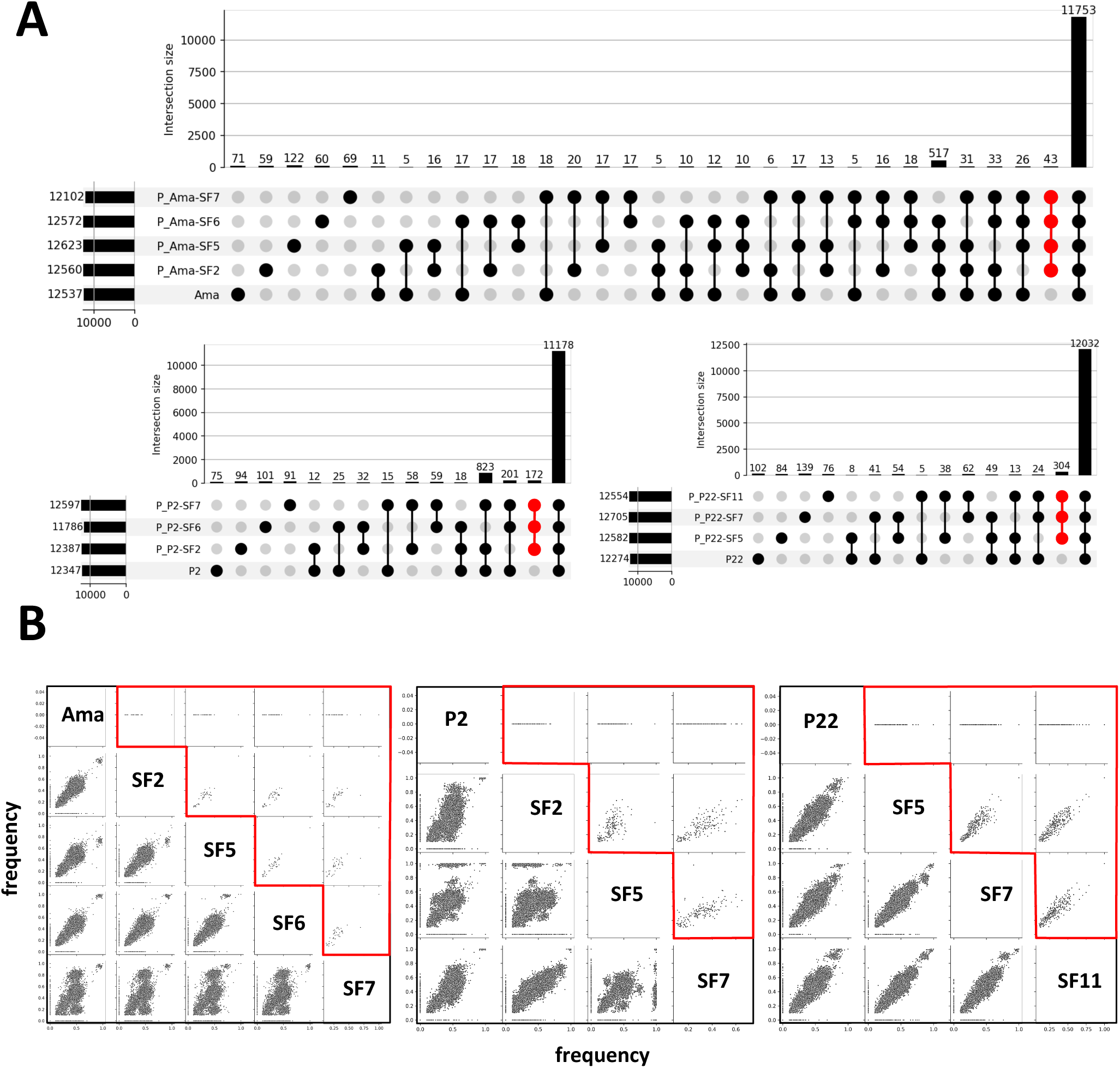
Genome-wide SNV analyses. (A) Upset plots of SNVs found in the different input parasites (Ama, P2, P22) and their corresponding output isolates. The bars visualize the number of SNVs common to a given combination of samples. In red are highlighted the SNVs that are common to only the output isolates. (B) Comparison of SNV frequencies between input parasites and their corresponding output isolates. Highlighted in red is the comparative analysis of SNVs that are common to the output isolates (as judged by the largely diagonal distribution pattern) and new compared to their respective input parasites.

Plotting SNV frequencies of the input parasites against their respective SF isolates revealed a series of interesting signals for the three infection groups (Figure 3B). Comparing the Ama input parasites with the P_Ama-SF2, 5 and 6 output isolates resulted in a prominent signal of SNVs between 10-70% frequency values, with an additional cluster observed at around 90%. The largely diagonal distribution of this comparison indicates that no major frequency shifts occurred in these SF isolates. In contrast, the comparison of Ama with P_Ama- SF7 generated a very different pattern, with most of the SNVs of the sand fly isolate showing frequency shifts to both higher and lower values. These shifts are neither explained by chromosome nor gene amplifications as judged by the largely constant read depth observed in this experimental group (see Figure 2). Likewise, comparison between the P2 input parasites and the P_P2-SF2 and P_P2-SF5 (and to a lesser extend also P_P2-SF7) isolates showed important SNV frequency changes independent of chromosome or gene read depth variations, which are visualized as patches of intermediate (0,6 – 0,75) or strongly increased frequency (> 0,75). Unlike the Ama and P2 infection groups, no significant differences were observed for the P22 input parasites and the derived P_P22-SF isolates as indicated by the diagonal SNV distribution. Comparison of only the SNVs that are new with respect to the parental input and shared by the SF isolates further sustains their convergent selection during sand fly infection (Figure 3B, red frames).

In conclusion, we observed important shifts in SNV frequency during sand fly infection that apparently occurred independent of chromosome or gene amplification. The most parsimonious explanation for these shifts is provided by transient changes in ploidy or genetic recombination events during sand fly infection – possibilities we further explored by a more in-depth analysis of the SNV signals.

### Haplotype selection occurs in the absence of stable aneuploidy during sand fly infection

We next analyzed the distribution of SNV frequencies, chromosome by chromosome, for the various samples (Figure 4). As expected from the presence of two haplotypes in *L. donovani* Ld1S [14] and the largely disomic state of the Ama input parasites, most heterozygous SNVs in the Ama sample are present at a frequency of 50% (or 0,5), giving rise to a single peak (Figure 4, upper left panel; see Figure S3 for the individual frequency distribution plots). The SNV frequency distribution for chr 31 shows a major peak at a frequency of 25%, which demonstrates that the four haplotypes of this tetrasomic chromosome are balanced (at least on population level). Chromosomes 4, 15 and 33 are strongly depleted in SNVs likely due to previous loss-of-heterozygosity events involving transient monosomies and subsequent chromosome duplication. While the Ama SNV profile is otherwise largely reproduced in the P_Ama-SF2, 5 and 6 isolates, the profile in P_Ama-SF7 shows important differences: First, this isolate shows a bimodal distribution for several chromosomes with peaks at 20% and 80% frequency (i.e. chr 1, 2, 14) indicating strong selection of one over the other haplotype. In the absence of any observed karyotypic change (see Figure 2A), such frequency shifts are likely explained by transient trisomy and strong haplotype selection as we previously reported [14]. Second, an even more complex SNV frequency distribution was observed for chr 3, 9, 10, 11, 21, 23, 25, 32, 35, and 36, that all maintain the 50% frequency peak but in addition show small peaks at low (<25%) and high (>75%) frequencies.

**Figure 4:**
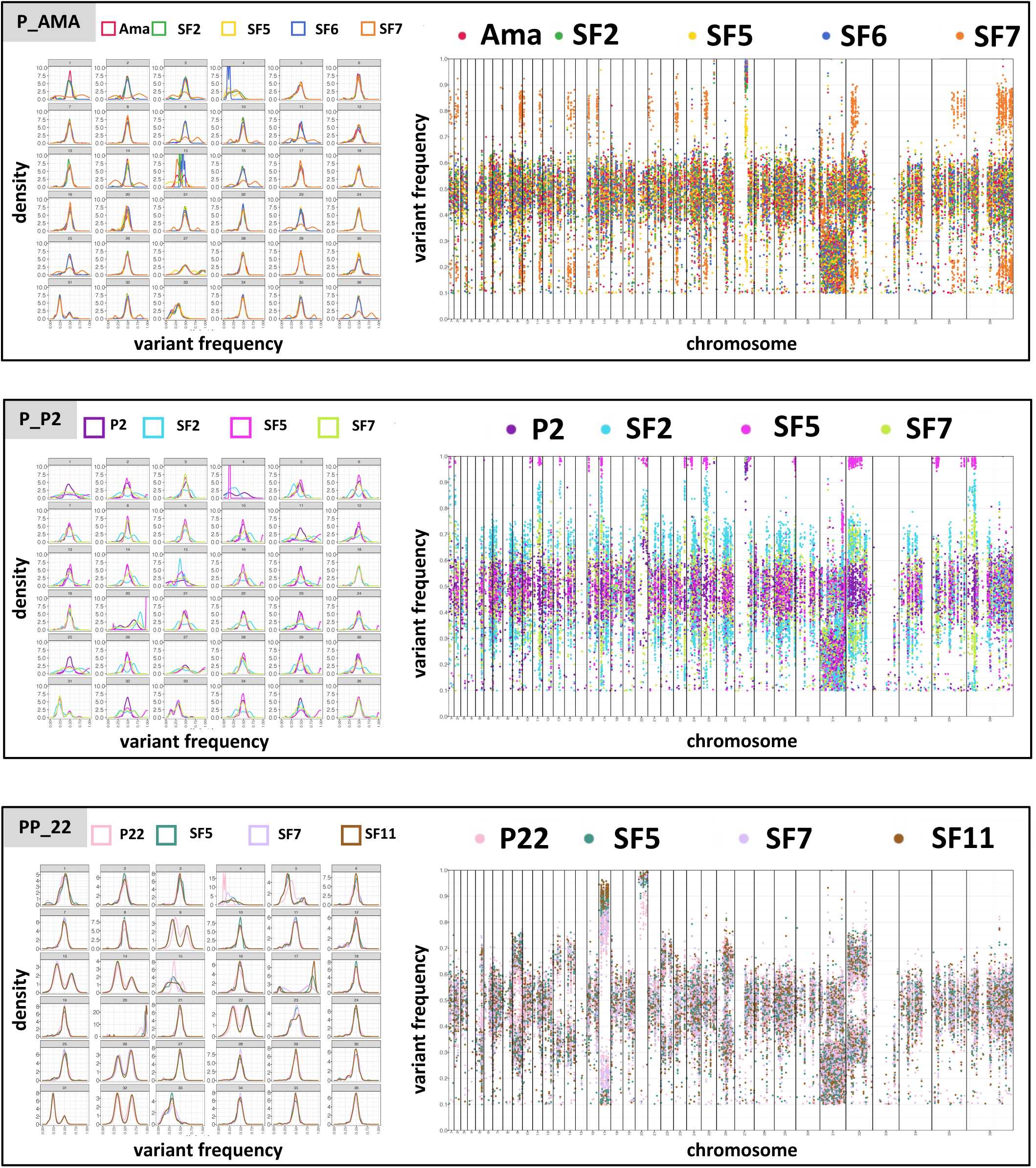
SNV frequency distribution. The plots on the left represent on the x-axis the variant allele frequency and on the y-axis the corresponding estimated probability density. Different chromosomes are reported in different panels. The scatterplots on the right display individual SNVs as dots. The x-axis indicates the genomic position while the y-axis indicates the variant allele frequency. The different chromosomes are displayed one after the other, and their boundaries are visualized as vertical lines. The lines (on the left) and the individual SNVs (on the right) are colored according to the samples.

P_P2-SF isolates showed more dynamic changes in the SNV pattern following sand fly infection compared to the P_Ama-SF isolates (Figure 4, middle left panel; see Figure S3 for the individual SNV frequency distribution). Many P_P2-SF chromosomes show a bimodal SNV frequency distribution not observed in the P2 parental parasites, with for example a 33/66% frequency ratio observed for various chromosomes of P_P2-SF2 (chr 2, 6, 8, 10, 14, 16, 17, 22, 26, 29, 34), which we previously associated with trisomy and haplotype selection [14]. The absence of aneuploidy in these P_P2-SF isolates (see Figure 2) further sustains transient trisomies as a possible mechanism allowing for haplotype selection in the sand fly midgut. This is well illustrated by chr 20 that is disomic in Ama showing a balanced haplotype distribution but is amplified in the Ama-derived P2 input parasites showing a 33/66% frequency distribution (Figures 2 and 4). All P_P2SF output isolates however are again disomic, now showing an extreme SNV frequency distribution of 10/90% that reveals the replacement of one haplotype by the other via transient aneuploidy. In addition, many chromosomes of P_P2-SF5 maintain a dominant peak at 50% SNV frequency, but show a new, low density peak at very high frequency (e.g. chr 2, 10, 16, 17, 23, 35, 36) that is absent in the P2 parental parasites.

In contrast to the Ama and P2 infection groups, the P_P22-SF isolates largely reproduce the profiles of their P22 parental parasites, which are characterized by a 33/66% bimodal SNV frequency distribution for chr 5, 9, 13, 14, 22, 26, 32 (Figure 4, lower left panel), which correlates with partial or full trisomies (Figure 2) and haplotype selection as a result of long-term culture adaptation [14]. Of note, the partial trisomies observed for chr 5 and 9 (see Figure 2) were further selected to full trisomies in all P_P22-SF isolates as indicated by the 33/66% frequency distribution (Figure 4, lower left panel; see Figure S3 for the individual SNV frequency distribution), suggesting that one of the haplotypes may provide an unknown fitness advantage to promastigotes proliferating in the sand fly midgut.

### Hot spots of sub-chromosomal SNV frequency shifts

We next investigated in more detail the sub-chromosomal changes in SNV frequency. Plotting variant frequency against chromosomal location (Figure 4, right panels, and Figure S4) revealed a patchy SNV distribution in all chromosomes of the input parasites Ama, P2, P22, with short zones of heterozygosity separated by zones devoid of SNVs. Similar to the Ama input sample, most P_Ama-SF isolates show a frequency distribution around 50% (Figure 4, upper right panel) as expected from their disomic state (Figure 2A) and the largely balanced distribution of the two haplotypes (Figure 4, upper left panel). In contrast, the isolate P_Ama-SF7 shows important shifts of individual SNV patches towards a 20/80% frequency distribution for chr 1, 2, 3, 7, 9, 10, 11, 14, 16, 21, 23, 25, 32, 35, 36, consistently with the bimodal and tri-modal SNV frequency profiles observed for these chromosomes. Likewise, two SNV patches on chr 31 show a frequency shift from 25% to 40% and 65% (see individual plots in Figure S4 for more detail).

We observed many sub-chromosomal SNV frequency shifts for all P_P2-SF isolates, with P_P2-SF5 attaining frequencies of close to 100% for SNV patches on chromosomes 1, 2, 10, 11, 13, 16, 17, 20, 23, 24, 25, 29, 32, 35, 36 (Figure 4, middle right panel). Significantly, for many of these SNV patches we also observed a significant increase in frequency in the other two SF isolates, ranging from 70% in P_P2-SF7 to 85% in P_P2-SF2. Given that these SF isolates are derived from different sand flies and thus can be considered independent biological replicates, this striking convergence both in SNV localization and frequency shift identifies hot spots of alleles that may be under positive selection during sand fly infection. In contrast to P_Ama- and P_P2-SF isolates, no major SNV frequency shifts were observed in the P_P22-SF isolates compared to the P22 input parasites, with the exception of SF7 that shows minor frequency shifts for chr 17 and 20 (Figure 4, lower right panel).

In conclusion, the observed SNV frequency shifts and their potential convergence in both location and frequency reveals an additional form of *Leishmania* genome instability during sand fly infection that may allow for haplotype shuffling and allelic selection.

### Haplotype shuffling during sand fly infection

We further investigated converging SNV frequency shifts by plotting the frequency difference between parental input and the SF output isolates. This analysis revealed that only a subset of SNVs undergo frequency shifts, some of which define hot spots that show frequency increase across all SF isolates in a given experimental group. The presence of such hot spots is visualized in Figure 5A for the two chromosomes 23 and 36 of the P2 experimental group (see Figures S5-S7 for the entire panel of all experimental groups). SNV frequency shifts largely involved short DNA regions and were not caused by copy number variation as judged by the constant read coverage between input and output parasites plotted in the same graph (Figure 5A).

**Figure 5:**
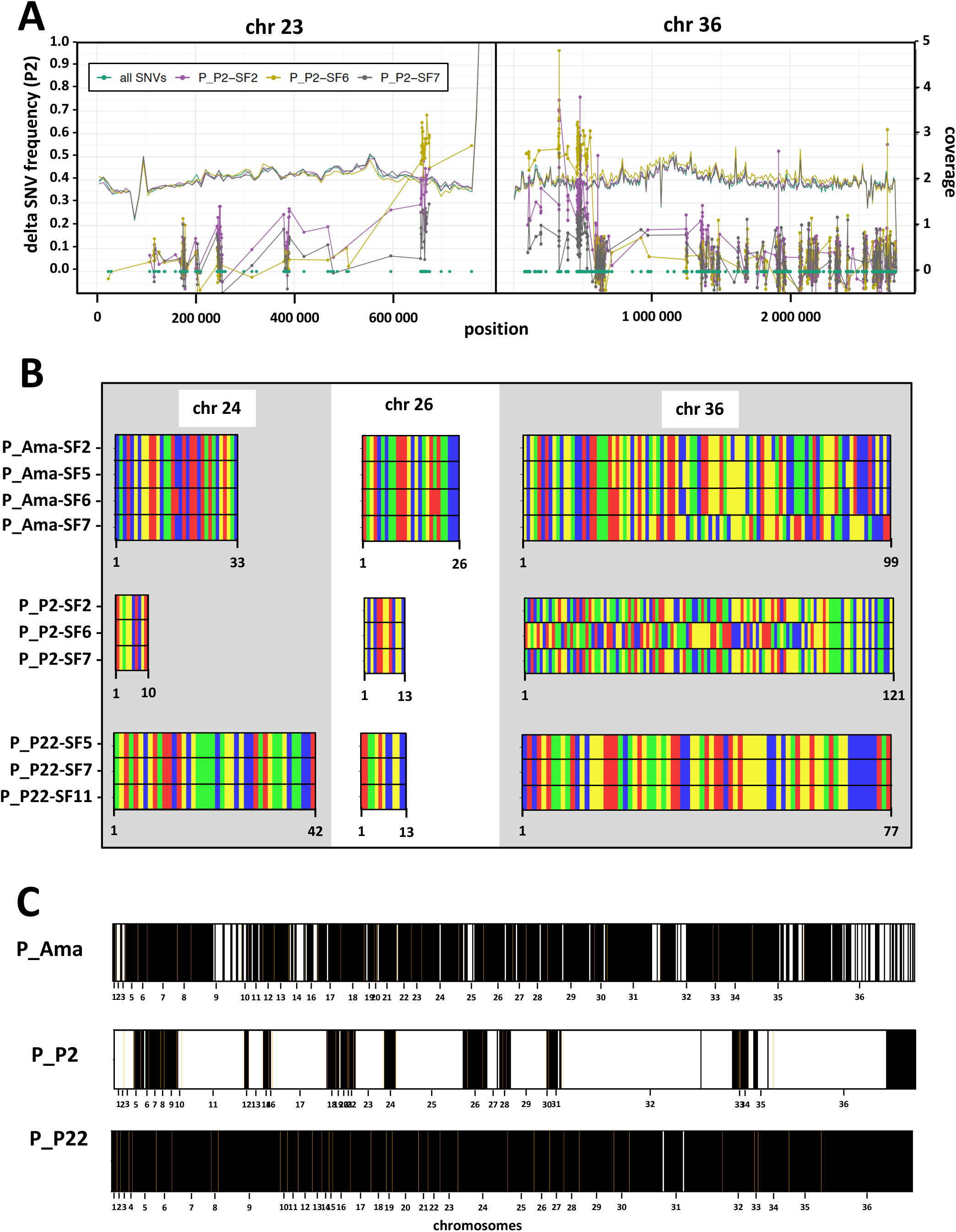
Convergent SNV frequency shifts and haplotype shuffling. (A) Differential SNV analysis. SNV frequency changes (“delta”) with respect to the input are represented for the output samples, along the chromosome coordinates. Positions of SNVs are represented as dots on the x-axis. For a given output sample, dots for successive SNVs are connected with lines, in order to better visualize possible “hotspots” for SNV frequency increases. The genomic coverage for a given output sample is indicated as a continuous line and is reported on the right y-axis. Only chromosomes 23 and 36 for samples corresponding to the P2 input are represented (see Fig. S5 – S7 for all panels). (B) Rainbow plot. SNV positions of the output samples that showed a strong frequency increase of either the reference allele or the alternate allele compared to their corresponding input samples are represented. A frequency increase is considered “strong” if it belongs to the top 5% percent (see Figure S8) in terms of “minimum absolute delta”, where the “minimum absolute delta” for a locus is the minimum across output samples of the absolute values of the allele frequency shifts with respect to the input. To visualize haplotype selection, the bars are colored according to the allele that shows the frequency increase (A: blue, C: green, G: yellow, T: red). Only chromosomes 24, 26 and 36 are represented. The number of SNVs per chromosome fulfilling our criteria is indicated below the x-axis. (C) Genome-wide assessment of haplotype selection. The rainbow plots for a given experimental group were scored for convergent (black bar) or divergent (white bar) alleles. Chromosome boundaries (orange line) and number are indicated.

We next investigated to what extent these frequency shifts affected the otherwise balanced haplotypes. As indicated above (see Figure 4) and published previously [14], the *L. donovani* Ld1S genome shows two distinct chromosomal haplotypes that are defined by heterozygous SNVs. The convergent SNV frequency shifts we observed therefore can be produced by both of the two alleles that represent a given heterozygous SNV. We next assessed if this choice occurs at random or if one allele is preferred over the other and thus may be under positive selection. Painting the dominant allele with distinct colors for the four possible nucleotide bases revealed a surprising degree of allelic selection, with many chromosomes showing a highly reproducible and converging SNV pattern either along stretches of a given chromosome (e.g. chr 36 of the P_Ama and P_P2 SF output parasites, Figure 5B) or the entire chromosomes (e.g. chr 24, 26, or 36 in the P_P22-SF output parasites). Plotting the allelic choice (i.e. selection of identical or different alleles between output isolates) for each convergent SNV position at genome-wide level shows that allelic selection affects all chromosomes in the Ama and P22 experimental groups, but only a subset of chromosomes in the P2 experimental group (Figure 5C).

In conclusion, our analyses reveal hot spots of SNV frequency shifts that generate new haplotype combinations, which may facilitate the selection of beneficial alleles during *Leishmania* sand fly infection.

### Allele frequency changes are associated with oxidative DNA damage

We investigated the potential biological impact of allelic selection by analyzing converging SNVs frequency shifts in coding sequence that caused non-synonymous mutations (Figure 6A-C and Tables S7-S9). For most of these examples, the parental input parasites show a zero-frequency value, indicating that either this SNV does not exist or has been filtered out by our computational pipeline due to quality issues linked to read mapping. Strikingly, these SNVs not only converged between the SF output isolates of a given experimental group, but some even between experimental groups. For example, the SNV 34_600750_T_G (defining chr_position_reference_variant) that causes an Asn221His amino acid (aa) change in a Rab-like GTPase activating protein on chromosome 34 (Ld1S_340628300) is observed in all P_Ama-SF and P_P2-SF isolates. Likewise, convergent SNVs between the P_P2 and P_P22 experimental groups have been observed for the ras-like protein Ld1S_360772200 on chr 36 (SNV 36_1917048_A_G, aa change Asn166Ser) and the hypothetical protein Ld1S_060842800 (SNV 6_494645_A_G, aa change Asn15Ser). How these mutations may affect protein structure, function and interactions, and if they establish beneficial parasite phenotypes remains to be elucidated. Converging frequency increases in the SF output isolates were also observed for synonymous SNVs that have no overt effect on gene function (see Tables S7-S9). Such synonymous SNVs may just be ‘hitchhiking’ with non-synonymous SNVs under selection. Alternatively, synonymous SNVs may alter the codon profile and thus may be selected for changes in translation efficiency, a possibility that is supported by our recent observation that *Leishmania* fitness gain correlates with tRNA gene amplification [10]. Finally, convergent SNVs were even observed in non-coding, intergenic regions (see Tables S7-S9). These mutations may again be ‘hitchhiking’ or could be selected for by beneficial changes in DNA topology or by changes in gene expression if affecting regulatory sequences in 3’ and 5’ UTRs that control RNA turn-over [36, 37].

**Figure 6:**
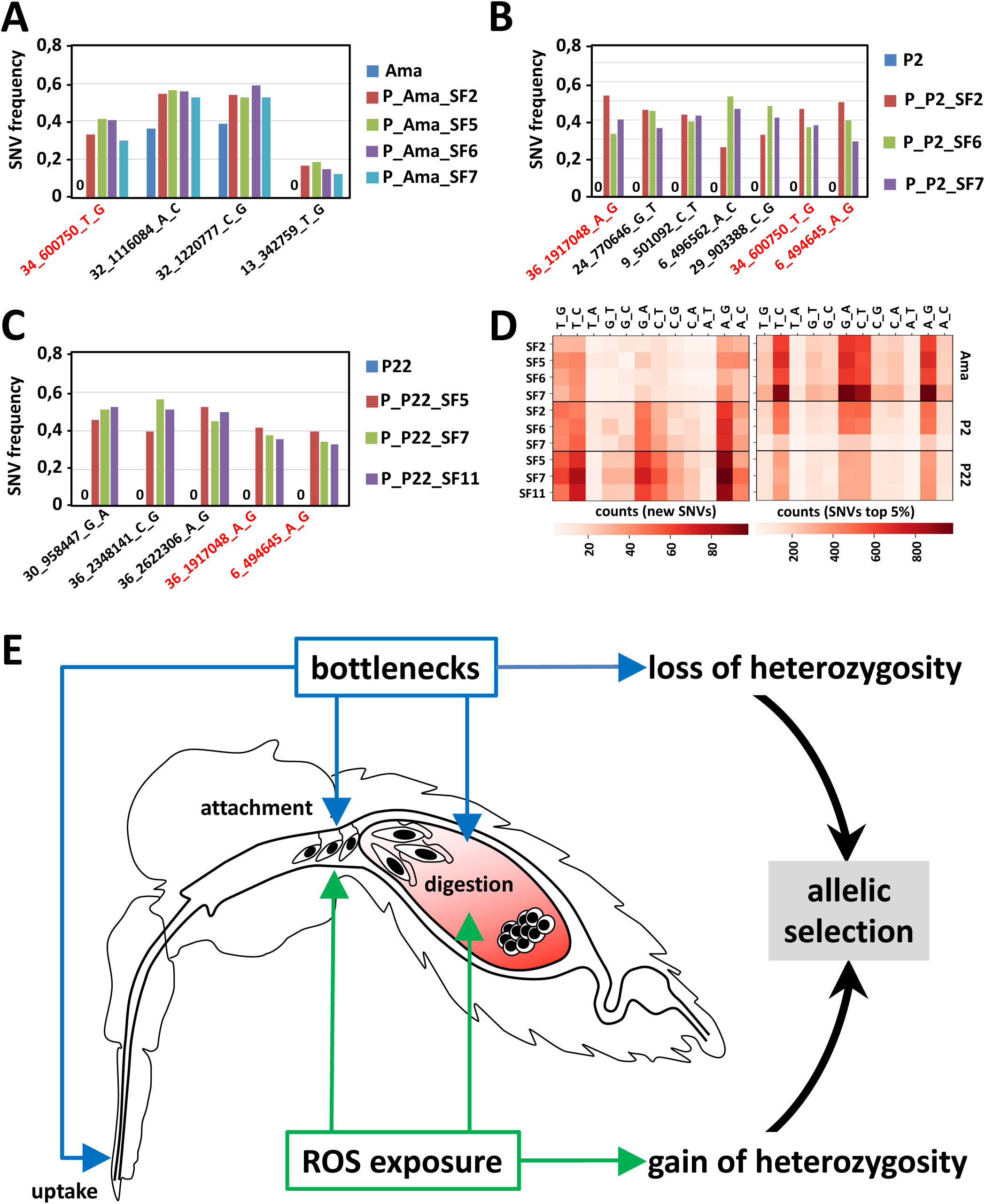
Allelic selection and oxidative DNA damage during sand fly infection. Plots showing the frequencies of the indicated, non-synonymous SNVs in the Ama (A), P2 (B), and P22 (C) experimental groups. SNVs that are shared between experimental groups are indicated in red. (D) Mutation distribution analysis. Heatmaps showing the distribution of the categories of mutations observed in the indicated output isolates compared to the corresponding input parasites. The color intensity indicates the count for the indicated mutation category according to the shown legend. The left heatmap represents the number of SNVs which are new in the sample with respect to its corresponding input. The right heatmap represents the number of SNVs whose abs_delta indicator was above the 95^th^ percentile in terms of min_abs_delta (see methods and Figure S8). (E) Model of *Leishmania* genomic adaptation during sand fly infection. Sand fly infection can affect the *Leishmania* population structure in two ways: (i) various bottlenecks (blue) that cause loss of heterozygosity in the parasite population due to random sampling during uptake, blood meal digestion, or parasites attachment to the midgut epithelium following excretion of the digested blood meal; (ii) ROS exposure and oxidative DNA damage that cause gain of heterozygosity through the generation of new SNVs. Both the bottleneck events and oxidative DNA damage contribute to allelic selection of potentially beneficial SNVs (see Discussion).

We next assessed the possible mechanism that triggers the observed allele frequency shifts. The observed mutations may be the result of parasite exposure to DNA-damaging agents, such as exogenous reactive oxygen species (ROS), known to regulate antimicrobial immunity and microbiome homeostasis in the insect gut [38], or endogenous ROS that can be generated in response to stress [39]. Oxidative DNA damage increases genome instability and introduces signature mutations due to mismatches caused by DNA repair intermediates [40]. To investigate signatures of ROS-mediated mutation, we quantified on genome-wide levels the type of nucleotide changes that caused allele frequency shifts in the SF output isolates compared to the corresponding input parasites. The induced nucleotide changes were highly reproducible for all SF isolates and involved mainly AóG and TóC transitions, which represent the majority of observed SNV changes (Figure 6D and Table S10). Such transitions are indeed diagnostic for oxidative DNA damage and can be caused by (i) oxidation of guanine to 7,8-dihydro-8- oxoguanine (8-oxo-G), (ii) oxidation of cytosine to 5-hydroxycytosine (5-OH-C) or cytosine to 5-hydroxyuracil (5-OH-U), and (iii) oxidation of adenine to hypoxanthine, all of which generate repair intermediates with altered base pairing properties [41]@.

In conclusion, our data establish a first association between DNA damage likely caused by exposure to oxidants that can generate genetic heterogeneity and allows for the selection of beneficial alleles during sand fly infection (Figure 6E).

## DISCUSSION

Here we combined our *Leishmania in vitro* evolution system [10, 14, 32] with experimental sand fly infection to assess the impact of the insect midgut environment on parasite genomic adaptation. We identified vector infection as an important genetic bottleneck that causes loss of heterozygosity and revealed haplotype shuffling and allelic selection that are likely driven by oxidative DNA damage that (Figure 6E). Our findings generate important new insight into mechanisms underlying *Leishmania* genetic diversity and evolvability, and open a series of questions on how *Leishmania* genomic adaptation drives parasite fitness inside its insect vector and what factors may cause oxidative DNA damage.

During their passage through their insect host, vector-borne pathogens (notably viruses) show a strong reduction in population heterogeneity, due to the limited number of microbes that are ingested or establish productive infection [42, 43]. In evolutionary biology, such reduction in population size due to environmental factors is referred to as a bottleneck, during which random sampling can have profound effects on the frequency of genotypes or alleles – a phenomenon known as genetic drift [44]. In contrast to this stochastic process, changes in allele frequencies can also be non-random and driven by natural selection for traits that are beneficial in a given environment [2]. Dissociating random from non-random allele frequency changes represents a key challenge in evolutionary studies. This holds true for our analysis on *Leishmania* genomic adaptation in the sand fly, which revealed a bottleneck event as judged by the polyphyletic distribution of input and output parasites, the non-reproducible karyotypic changes and the loss of SNVs following sand fly passage, indicating that certain parasite sub- populations were eliminated from the infecting population. This bottleneck is likely caused by (i) the small number of parasites ingested during a sand fly blood meal, (ii) the potential elimination of parasites by toxic byproducts during blood meal digestion, or (iii) their failure to attach to the midgut that leads to parasites excretion via defecation of the digested blood meal [29, 34, 45–48] (Figure 6E).

Despite random genomic changes caused by these bottlenecks, the comparison of independent biological replicates allowed us to also reveal non-random and thus selected genomic signals on chromosome, gene and especially nucleotide levels. We identified several SNV frequency shifts that converged not only between the biological replicates inside, but even between experimental groups, providing strongest support for natural selection. Even though the phenotype associated with these mutations eludes us, the annotation of the respective genes allows to speculate on the mechanism of fitness gain. For example, two of the converging alleles affected distinct members of the Ras GTPase family and thus a signaling pathway known to regulate eukaryotic cell growth, division and differentiation [49]. Indeed, Ras GTPases have been linked to cell proliferation and differentiation in *Trypanosoma cruzi* [50, 51], microtubule biogenesis and cytokinesis in *T. brucei* [52] and their survival during tse tse fly infection [53]. Significantly, in *L. donovani*, a non-synonymous SNV in the Ras-like small GTPase-RagC (LdBPK_366140.1) affects tissue tropism [15], indicating that a single SNV may produce important phenotypic change during infection. Hence, the convergent amino acid changes we have observed during sand fly infection in *L. donovani* Ras GTPases may drive fitness gain by increasing the parasite’s reproductive capacity inside the vector gut, for example by enhancing proliferation or promoting parasite attachment, possibilities that need to be investigated in future studies for example using CRISPR/Cas9 gene edited parasites.

The sand fly environment seems surprisingly conducive for *Leishmania* allele frequency changes and allelic selection considering the short, 8-day sand fly infection period. This could be a consequence of the bottleneck effect as the reduction of population heterogeneity allows for fast penetration of rare variants, even if they have only a minor beneficial effect [54]. On the other hand, such loss of heterozygosity has been linked to reduced evolvability [55], which could prove fatal once these parasites encounter subsequent bottle necks during transmission to the vertebrate host. The observed SNV dynamics thus call for an active mechanism that can replenish *Leishmania* genetic heterogeneity and fuel natural selection. We propose oxidative DNA damage as such a key mechanism as judged by the highly homogeneous, diagnostic mutation signature we observed across all analyzed output genomes, i.e. transitions between AóG and TóC [40, 41].

Oxidative DNA damage can be caused by reactive oxygen species (ROS), such as superoxide anion (O_2_^−^), hydroxyl radical (OH^−^) and hydrogen peroxide (H_2_O_2_) [40], which are a constant threat to DNA integrity and have been identified as a key driver of genome instability in various systems [40, 56]. ROS can be generated from various external and internal sources. In the context of *Leishmania* sand fly infection, ROS may be produced intrinsically inside the parasite as an enzymatic byproduct of the amastigote-to-promastigote differentiation process that is linked to important structural changes and retooling of parasite metabolism [57]. Likewise, the transition from an anaerobic, intracellular lifestyle (amastigotes) to an aerobic extracellular lifestyle (promastigotes) is associated with a shift in energy production that may result in ROS production [58].

On the other hand, ROS may be produced by external sources inside the sand fly gut. Indeed, ROS were shown in *Drosophila* to participate in the immune protection to pathogenic bacteria [59–61], and are produced in *Anopheles* (*A.*) *gambiae* and in *A. aquasalis* following infection with the parasite *Plasmodium falciparum* [62] and *Plasmodium vivax* [63], respectively. Even though *Leishmania* infection itself seems not to elicit detectable ROS inside the infected sand fly gut [64] a possible role of base-level ROS in affecting *Leishmania* genome integrity is supported by indirect evidence. First, the sand fly genome encodes for a member of the NAPDH oxidase family responsible for the generation of ROS (termed dual oxidase, DuOx) and several ROS detoxification enzymes, including superoxide dismutase and catalase (https://vectorbase.org/vectorbase/app/record/dataset/TMPTX_llonJacobina). Second, sand flies are able to produce immune protective ROS as shown in *Lutzomyia (L.) longipalpis* orally infected with the pathogenic bacteria *Serratia marcescens* [64]. Finally, the same study showed that silencing of the antioxidant enzyme catalase in *L. longipalpis* flies and the oral administration of ROS to the infected flies lead to a decreased number of *L. mexicana* parasites in the gut. The mutation signature observed in our experimental system lends further support to the exposure of *Leishmania* to ROS inside the sand fly gut. Whether such external ROS is directly produced by the insect gut cells, or indirectly by the endosymbiont bacteria composing the microbiota or toxic byproducts of bloodmeal digestion and hemoglobin catabolism [47, 65, 66] remains to be assessed.

Independent of the source of toxic oxidants, it seems that *Leishmania* may exploit oxidative DNA damage to promote its evolvability and increase its adaptive phenotypic landscape. However, such a strategy bears the risk to accumulate irreversible and damaging mutations, which is especially the case in asexual populations. This phenomenon is known as ‘Muller’s Ratchet’, which can only be alleviated by sex or recombination to allow efficient restoration of the wild-type sequence [67, 68]. As a matter of fact, *Leishmania* shows a cryptic sexual cycle capable of producing hybrid genotypes during sand fly infection, which can be induced *in vitro* by culture exposure to ROS [69]. *Leishmania* ROS-induced hybridization thus counteracts ‘Muller’s Ratchet’, while at the same time further increasing genetic variability and evolvability.

In conclusion, our results suggest that *Leishmania* exploits the DNA damaging environment in the sand fly midgut to increase its genetic heterogeneity and thus to further expand the already vast adaptive landscape of these parasites defined by other forms of genome instability, including chromosome and gene copy number variations [10, 14, 32]. Our study sets the stage for future investigations that aim to assess the quantity and quality of DNA damage during sand fly infection using single cell sequencing technologies, and to analyze the impact of ROS-generating and -detoxifying pathways on *Leishmania* genomic adaptation during sand fly infection by applying CRISPR/Cas9-mediated gene editing on both the vector and the parasite [69, 70].

## Supporting information

Table S1: Accession numbers for genome data

Table S2: Read mapping metrics

Table S3: AMA karyotype analysis

Table S4: P2 karyotype analysis

Table S5: p22 karyotype analysis

Table S6: Gene CNV analysis

Table S7: Ama_all_best_min_delta_above_0.12

Table S8: P2_all_best_min_delta_above_0.2

Table S9: P22_all_best_min_delta_above_0.3

Table S10: Mutation count frequencies

## Supplementary figure legends

**Figure S1: Genomic read depth variations.** Dots demonstrate the genomic bins. The x-axis indicates the normalized genomic bin sequencing coverage values. The y-axis reports the genomic position. To ease visualization, coverage values greater than 5 are reported as 5. The different rows and columns demonstrate the genomic coverage in different chromosomes and samples, respectively.

**Figure S2: Genome-wide SNV analyses.** Venn diagram (A) and SNV frequency comparison (B) of Ama, P2 and P22 input strains. (C) Venn diagram of SNV distribution in the input parasites and corresponding output isolates. Red numbers correspond to SNVs absent in the input and present in all outputs.

**Figure S3: SNV frequency distribution plots.** The x and y axes show respectively the variant allele frequency and the kernel density estimate. VRF, Variant Read Frequency (synonymous to variant allele frequency).

**Figure S4: SNV genomic localization.** The dots indicate SNVs and they are coloured according to the sample. Each panel represents a chromosome. The x-axis reports the genomic position. The y-axis shows the variant frequency.

**Figure S5: SNV localization, frequency shifts and read depth variation for the Ama experimental group.** SNV frequency changes (“delta”) in the output isolates with respect to the input parasites are represented along the chromosome coordinates. Positions of SNVs are represented as dots and reported on the x-axis. For a given output sample, dots for successive SNVs are connected with lines, in order to visualize possible “hotspots” for SNV frequency increases. The horizontal line indicates the genomic coverage for a given output sample as inducted by the second y-axis (right).

**Figure S6: SNV localization, frequency shifts and read depth variation for the P2 experimental group.** See legend of Figure S5 for details.

**Figure S7: SNV localization, frequency shifts and read depth variation for the P22 experimental group.** See legend of Figure S5 for details.

**Figure S8: Distribution of the min_abs_delta indicator across loci in the three experiments.** Left panels show the whole distribution while right panels show a zoom on the top 5% of the distribution. Different possible thresholds were considered to serve a loci selection criteria: 95^th^ percentile, 99^th^ percentile and mean plus two standard deviations. Their rounded values are represented using vertical lines, and the number of loci above these thresholds are indicated in the corresponding legend entry. In each histogram bin, the number of loci falling within an annotated gene is further highlighted in orange, and among these, the number corresponding to loci where the frequency shift with respect to the input follows the same trend is further highlighted in green.

## Acknowledgements

We thank Lucy Glover and Shaden Kamhawi for helpful discussions. This study was supported by a seeding grant from the Institut Pasteur International Department to the LeiSHield Consortium and the EU H2020 project LeiSHield-MATI - REP-778298-1. Prague group is supported by European Regional Development Fund (project CZ.02.1.01/0.0/0.0/16_019/0000759). Isabelle Louradour is financed by a Roux-Cantarini fellowship (S-FB14002-74A).

## Notes

https://www.ncbi.nlm.nih.gov/biosample/SAMN07430226

### Competing Interest Statement

The authors have declared no competing interest.

