## Supplementary figures and images for "*Leishmania* allelic selection during experimental sand fly infection correlates with mutational signatures of oxidative DNA damage"

### supplementary figure file

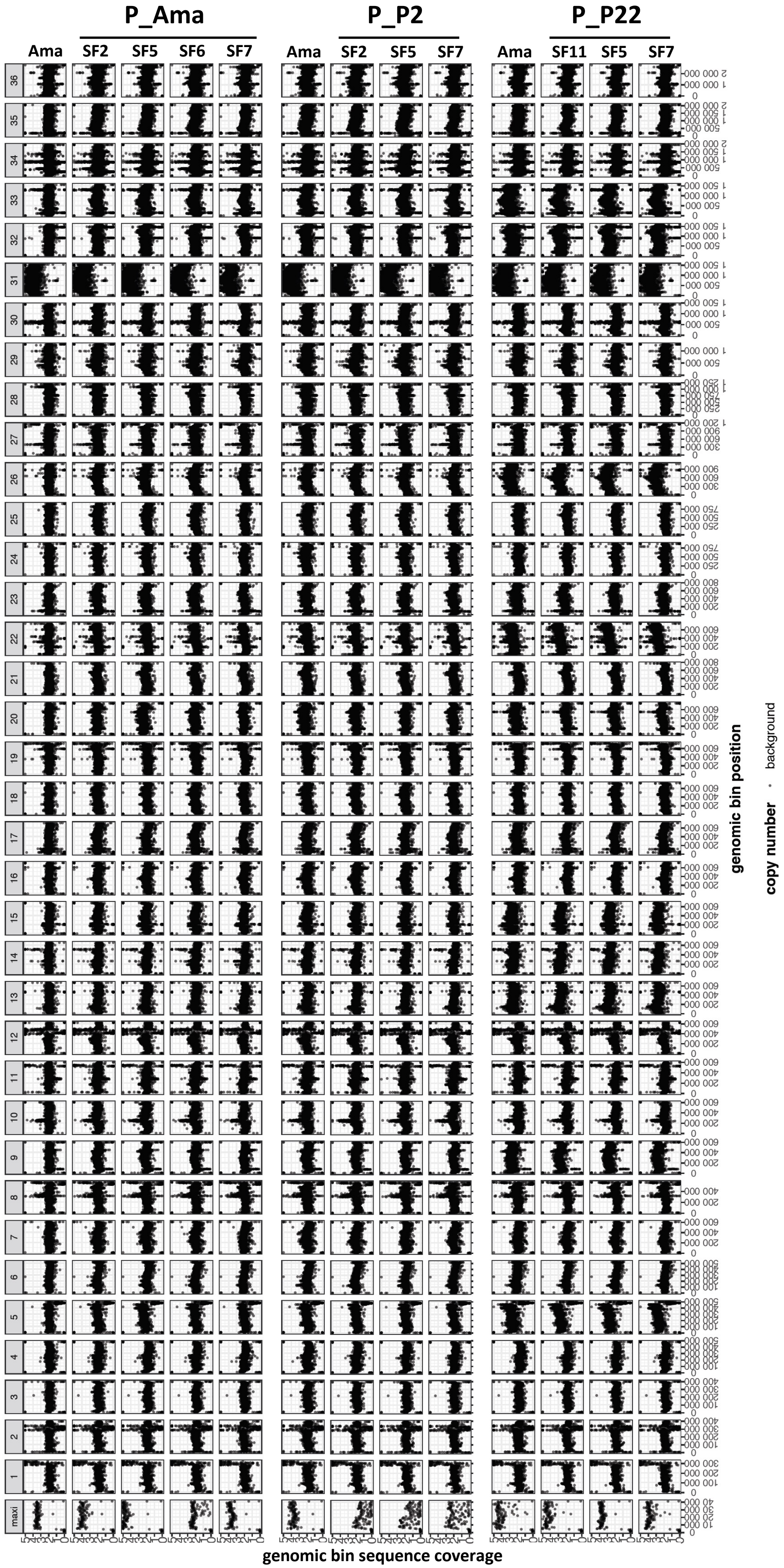

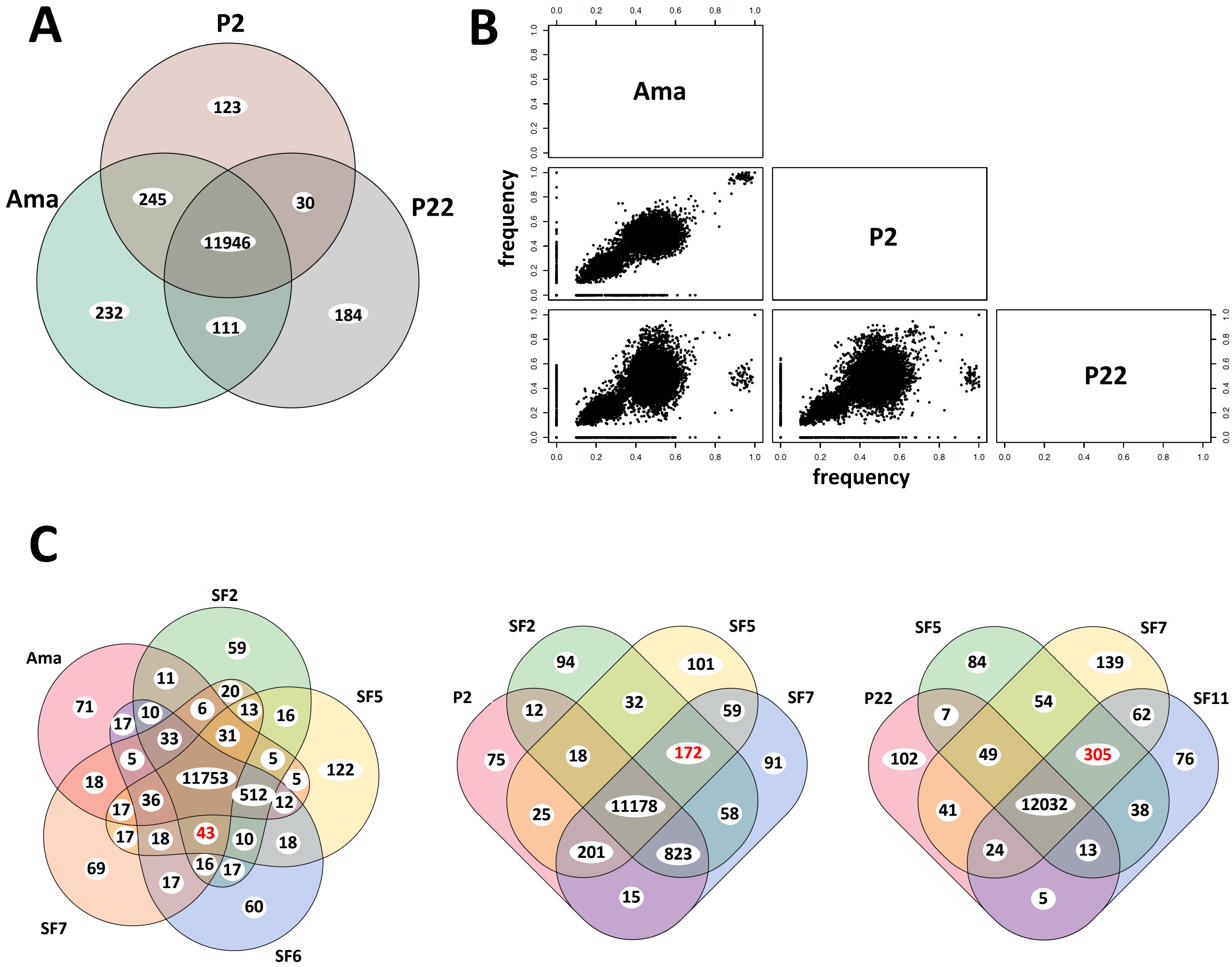

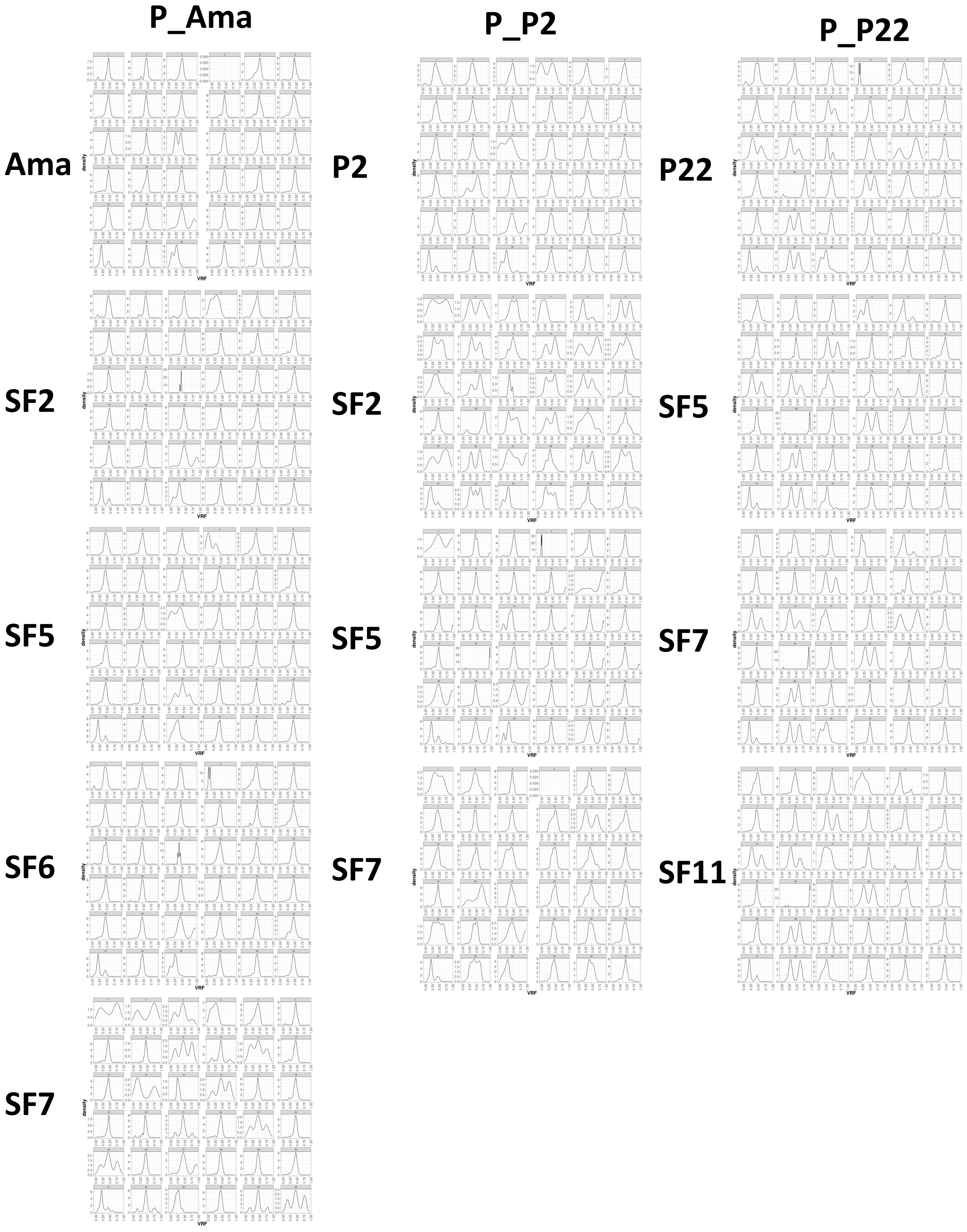

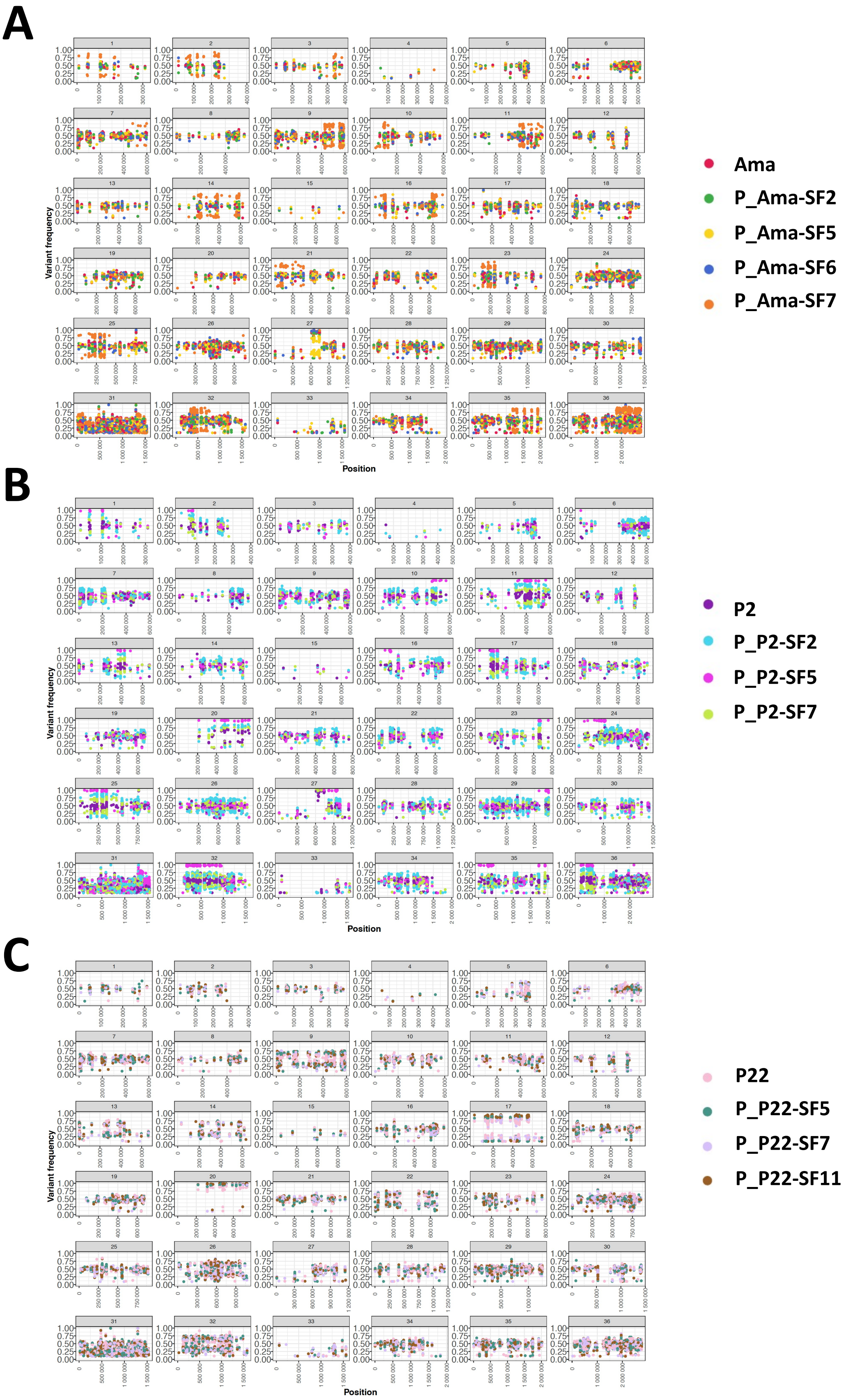

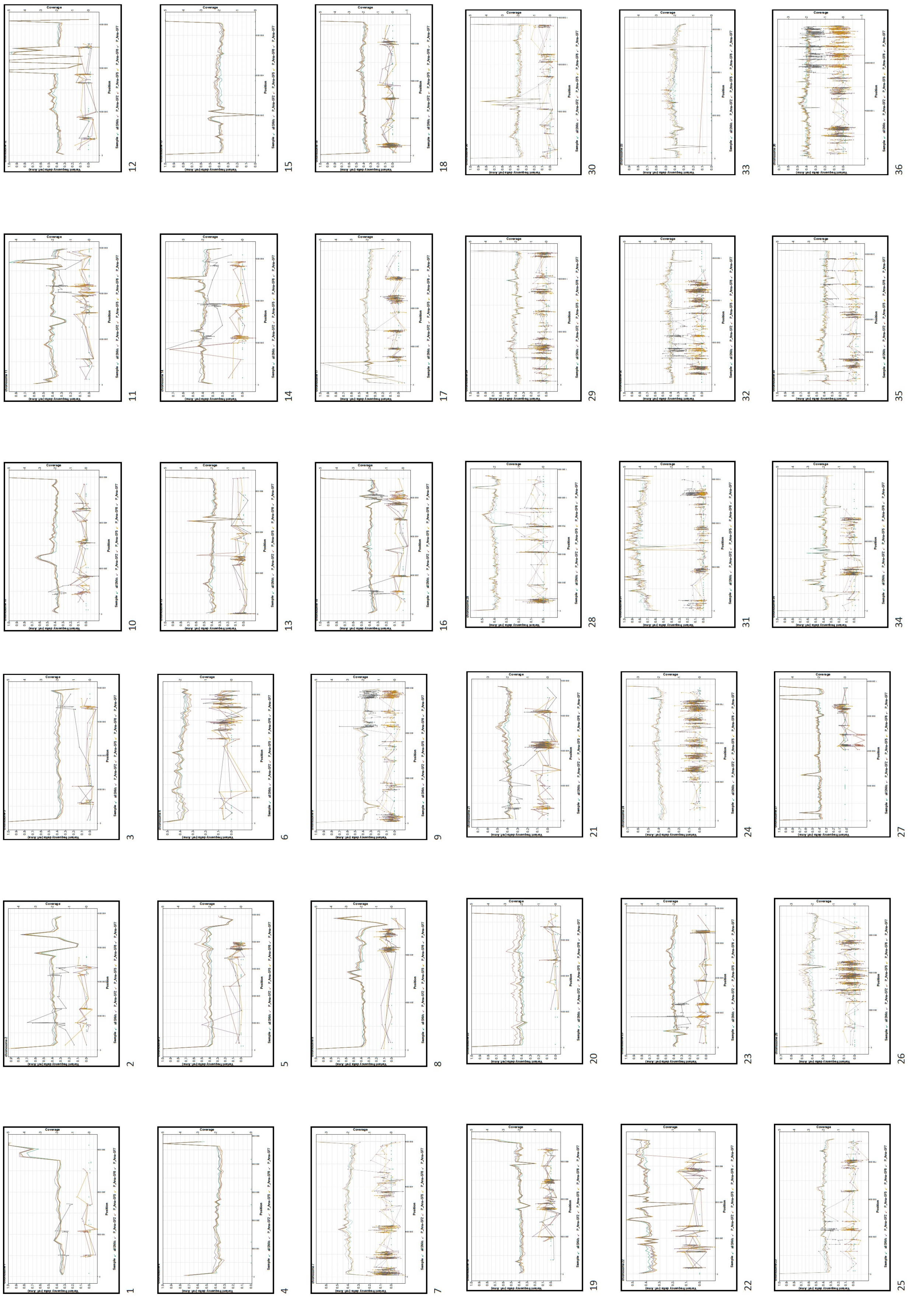

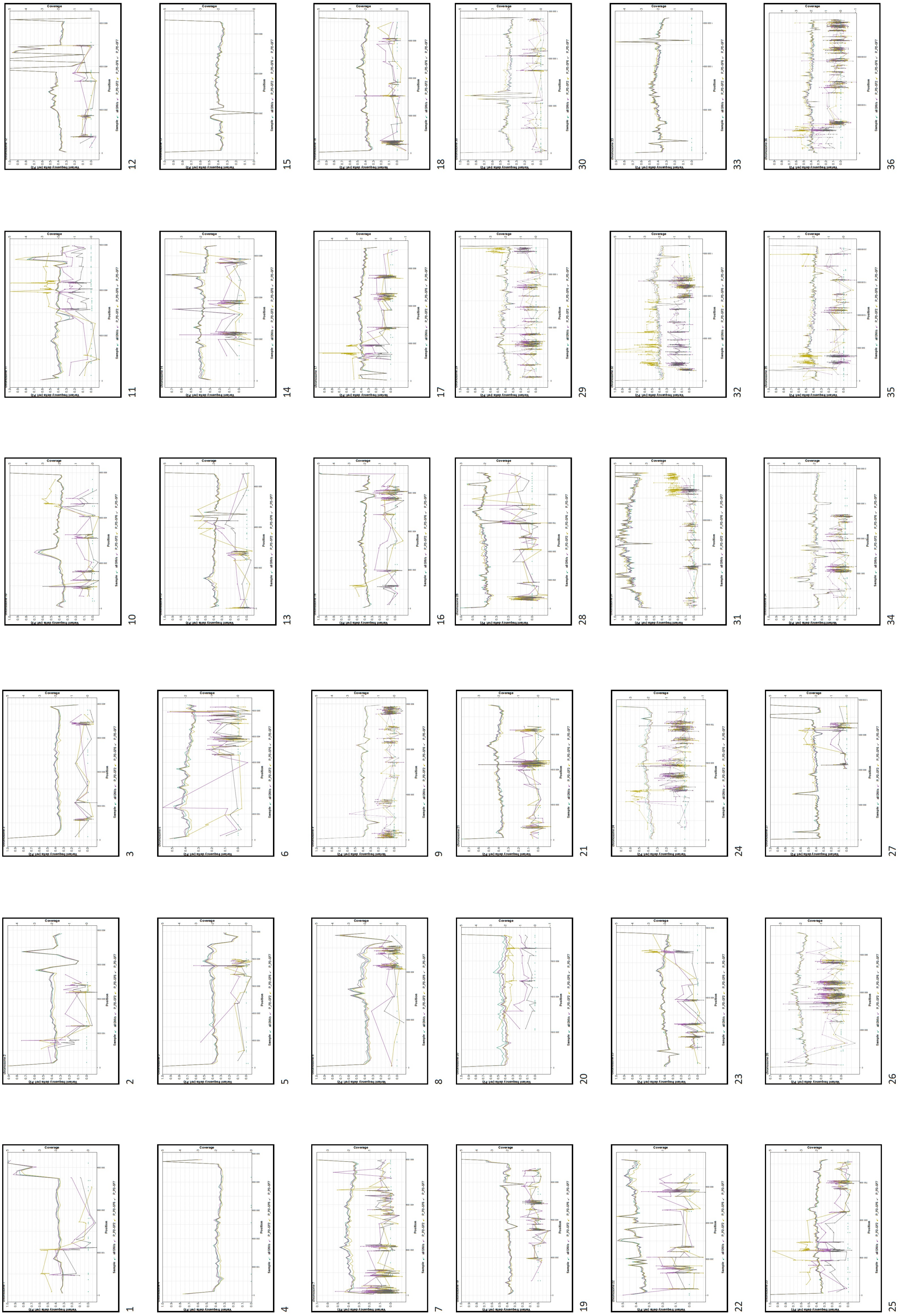

**Figure S6**

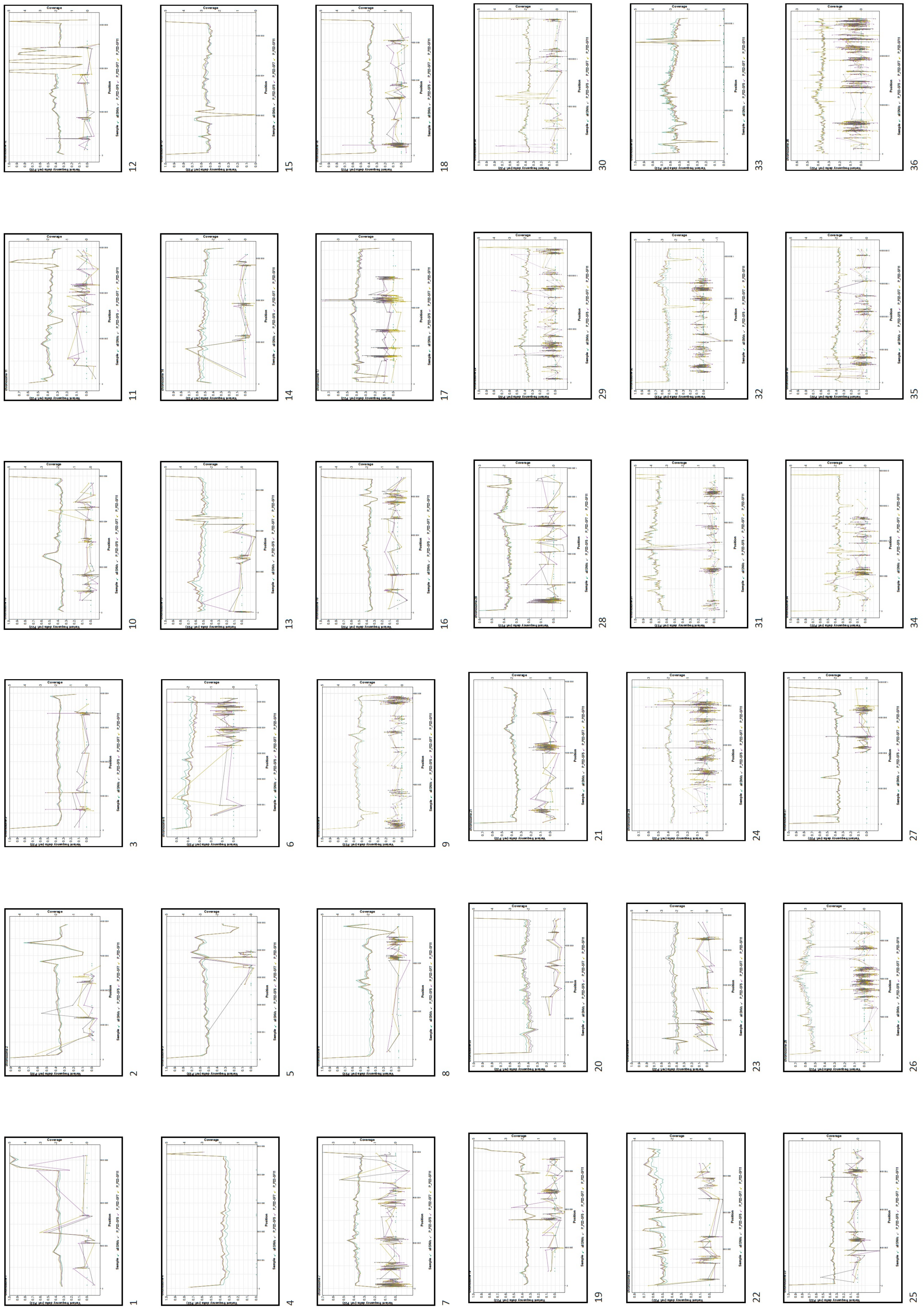

**Figure S7**

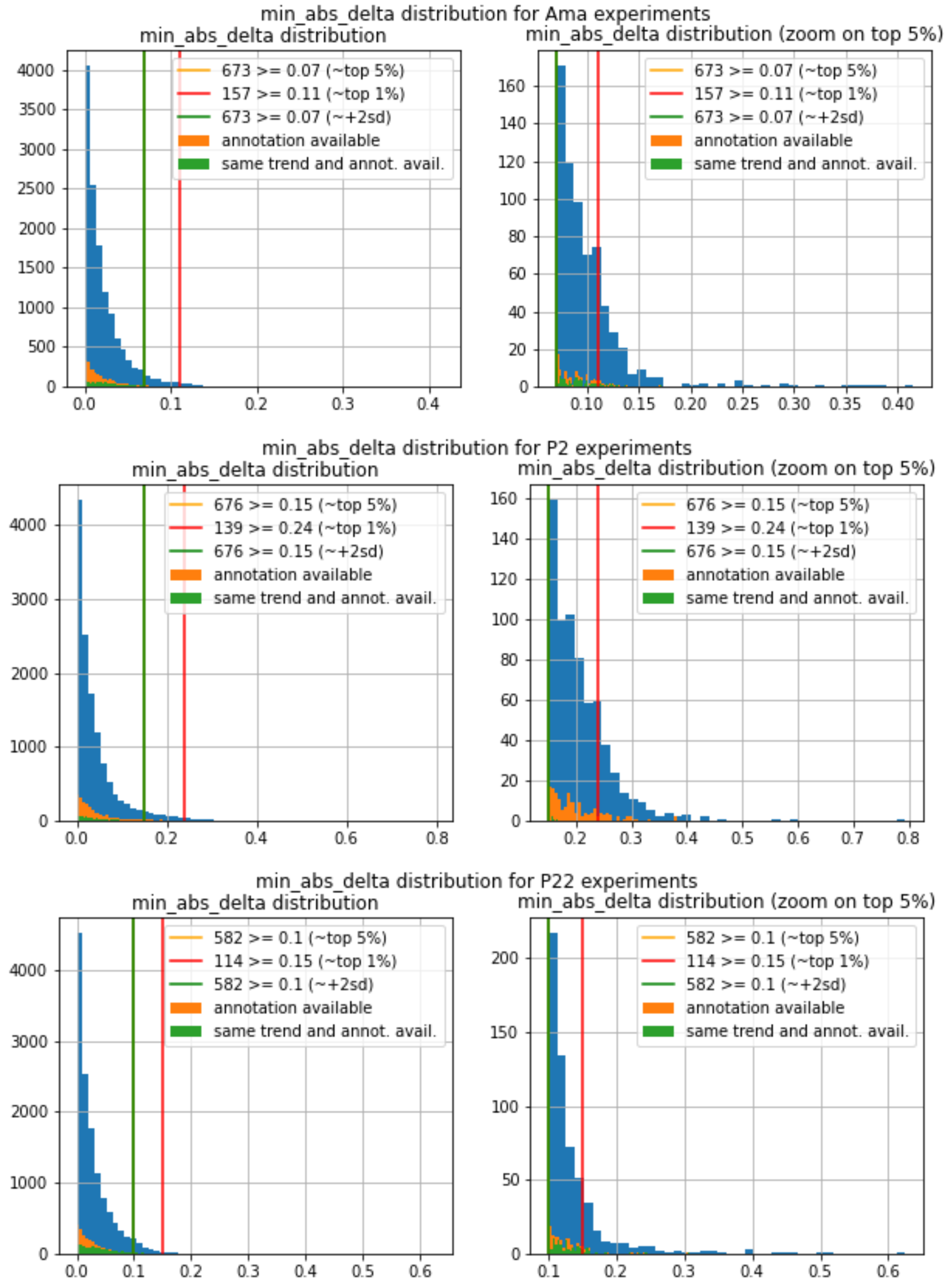
